# Development of the duodenal, ileal, jejunal and caecal microbiota in chickens

**DOI:** 10.1101/768747

**Authors:** Laura Glendinning, Kellie A Watson, Mick Watson

## Abstract

**Background:** The chicken intestinal microbiota plays a large role in chicken health and productivity and a greater understanding of its development may lead to interventions to improve chicken nutrition, disease resistance and welfare.

**Results:** In this study we examine the duodenal, jejunal, ileal and caecal microbiota of chickens from day of hatch to 5 weeks of age (day 1, 3, 7, 14 and week 5). DNA was extracted from intestinal content samples and the V4 region of the 16S rRNA gene was amplified and sequenced. We identify significant differences in microbial community composition, diversity and richness between samples taken from different locations within the chicken intestinal tract. We also characterise the development of the microbiota at each intestinal site over time.

**Conclusions:** Our study builds upon existing literature to further characterise the development of the chicken intestinal microbiota.

## Background

Improvements in sequencing technologies have led to a better understanding of the microbiota of many livestock species [1–4], leading some to suggest that we could optimise the composition of microbiota in these economically important animals to improve production and sustainability [5]. However, to do so would require a good understanding of the types of microbes which naturally occur in these animals and the role they play in nutrition and health.

Several studies have used 16S rRNA gene data to characterise the microbial communities which colonise the gastrointestinal tracts of chickens and to characterise the development of these communities over time. The vast majority of these studies have focussed on the chicken caeca as this is where the largest concentration of microbes can be found. The caecal microbiota has been suggested to play an important role in nutrition via the production of short chain fatty acids, nitrogen recycling and amino acid production [6–8]. In early life it is generally observed that the caeca contain high abundances of *Enterobacteriales* [9, 10] and over the first few weeks of life these decline and members of the *Clostridiales* come to predominate [9–16], with some studies also showing a large increase in *Bacteroidetes* [12, 17, 18]. However, the results from some studies do not entirely follow this pattern [19, 20] and variability in microbiota composition between flocks can be high [21, 22]. Several studies have also examined samples from the small intestine which are less rich and diverse than caecal samples and contain a high abundance of Lactobacilli [12, 14, 17, 18, 20, 23–25].

Several studies have directly compared samples taken from the small intestine with those from the caeca at specific timepoints [17, 18], and at various life stages. Lu *et al.* compared ileal and caecal samples from commercial Ross-hybrids, taking samples at day 1, 3, 7 14, 21, 28, and 49 days of age [14]. A separate study examined ileal and caecal samples from Hy-Line W-36 commercial layers at 9 timepoints, starting at 1 week of age [20]. Ileal and caecal samples were also characterised at various timepoints starting at 1 week of age by Johnson *et al.* in Cobb 500 birds [23].

In this study we compare the bacterial microbiota of duodenal, jejunal, ileal and caecal samples of Ross 308 broilers at 5 timepoints: 1 day, 3 days, 7 days, 14 days and 5 weeks of age. Similar to the studies above, we find significant differences in the microbiota compositions of specific sample types with age and significant differences between sample types within timepoints.

## Results

The V4 region of the 16S rRNA gene was amplified and sequenced, producing a total of 11,115,696 paired-end reads from 164 samples. 59.9% of sequences were removed during quality control. For each sample the average number of reads after quality control was 27,184 ± 44,159 (mean ± standard deviation). A total of 6015 operational taxonomic units (OTUs) were identified. **Additional file 1** contains the complete OTU table and **Additional file 2** contains the taxonomic assignment of OTUs. The most abundant bacteria in our negative controls were *Clostridium_sensu_stricto_1* (0.23 ± 0.20), *Lactobacillus* (0.17 ± 0.12), unclassified members of the *Lachnospiraceae* (0.14 ± 0.19) and *Enterococcus* (0.10 ± 0.09). The composition of the mock community control can be found in **Additional file 3: tables 1 and 2**. Prior to analysis, samples were subsampled to 10,000 reads; 35 samples were discarded as they had less than 10,000 reads, including all reagent only controls and 11 duodenum samples, 5 ileum samples and 4 jejunum samples from day 1. 2 jejunum samples from 5 weeks were also discarded. This resulted in only one duodenum sample from the day 1 timepoint remaining. The lowest Good’s coverage value for any of the remaining samples was 0.985, meaning that at least 98.5% of the bacteria in these samples were identified. Samples were compared statistically to see if there were differences in the microbiota between sample types at specific timepoints and if there were changes in specific sample types at different timepoints (**Figures 1 and 2**). The number of OTUs shared between sample types within timepoints and between timepoints within sample types can be found in **Additional file 3: figures 1 and 2**. Richness and diversity measurements for each subsampled sample can be found in **Additional file 3: table 3.**

**Figure 1:**
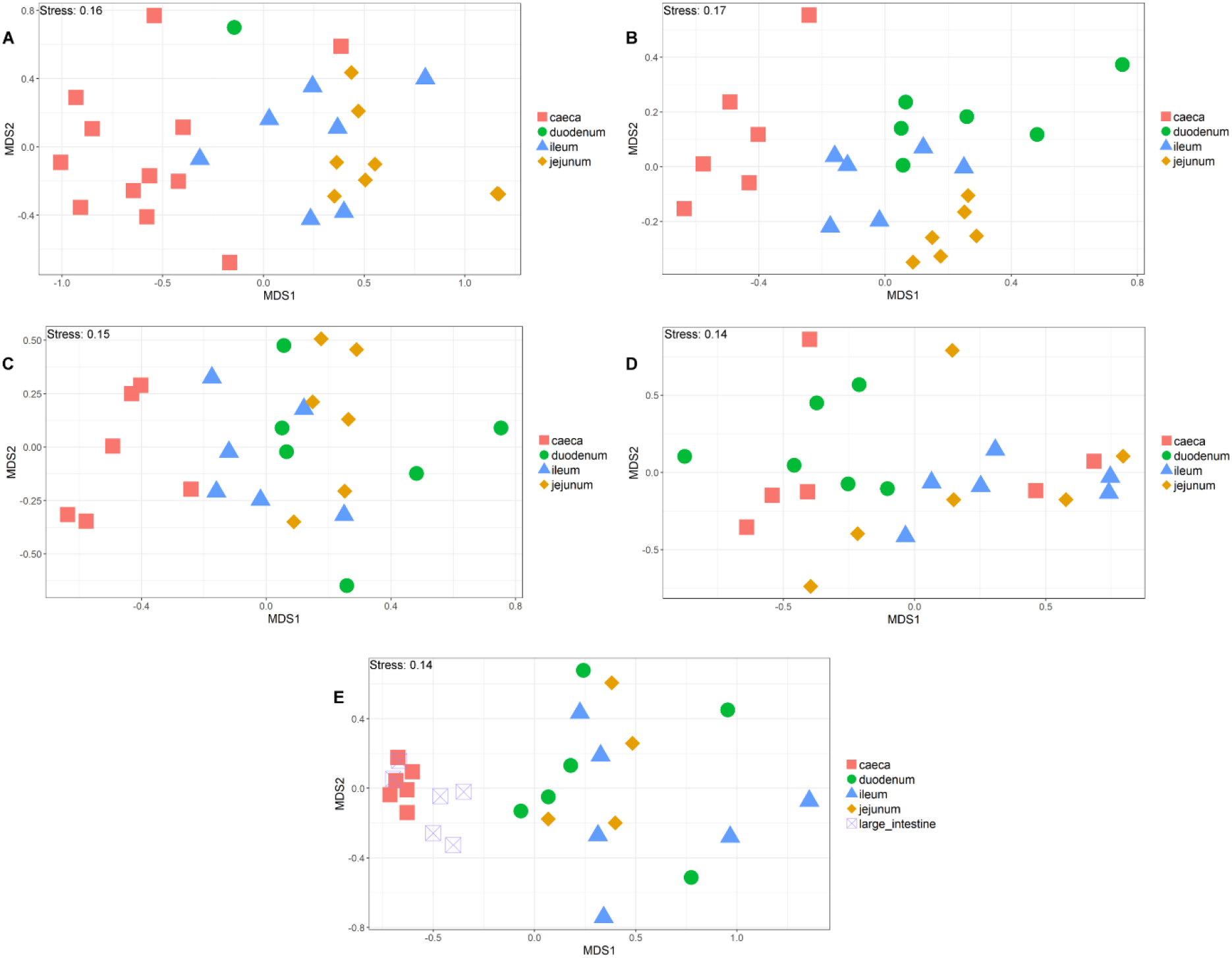
NMDS comparing sample types within timepoints. NMDS of chicken intestinal content samples clustered by microbial community composition (Bray-Curtis dissimilarity). A: Day 1, B: Day 3, C: Day 7, D: Day 14 and E: 5 weeks.

**Figure 2:**
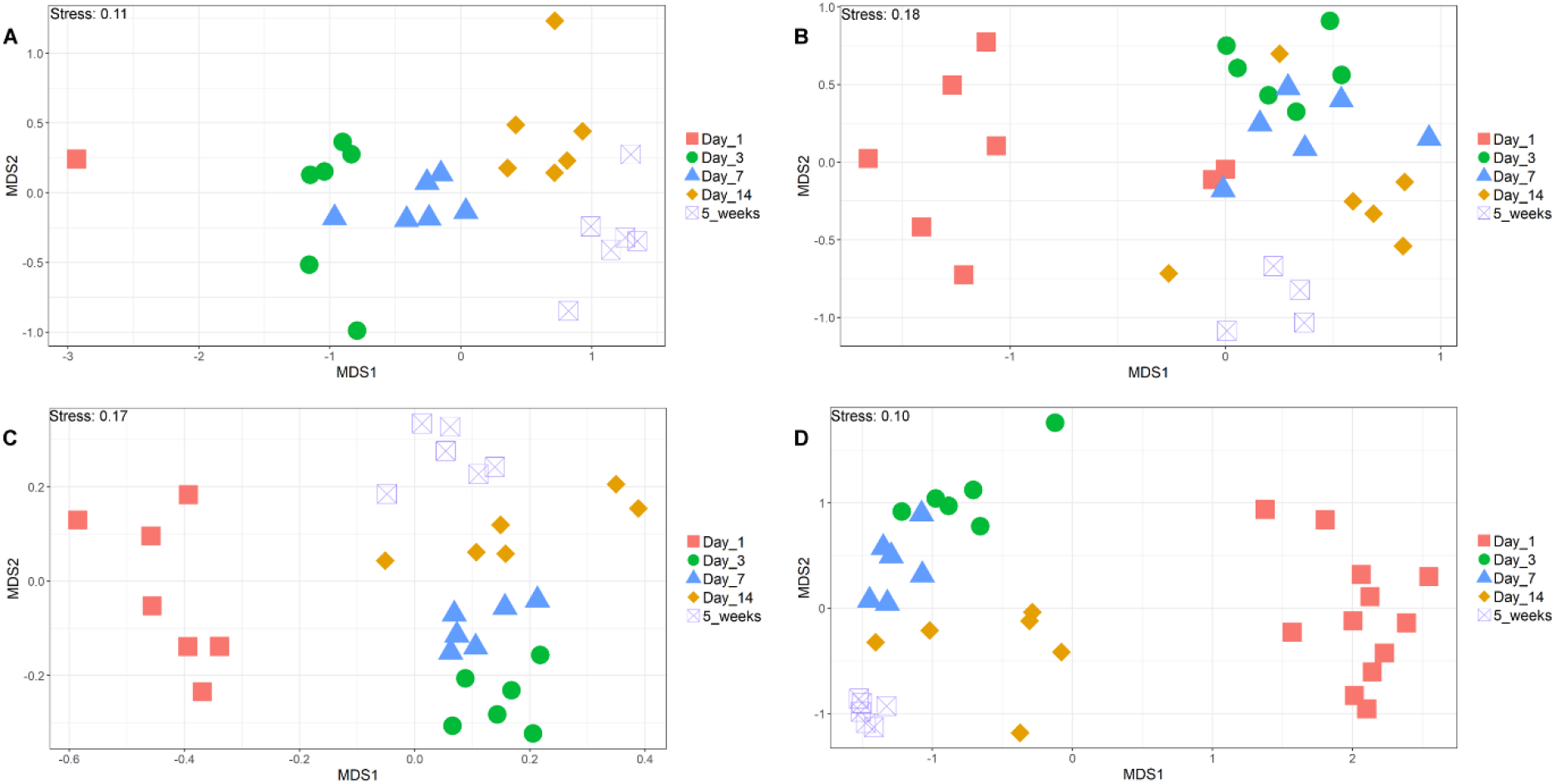
NMDS comparing sample types across timepoints. NMDS of chicken intestinal content samples clustered by microbial community composition (Bray-Curtis dissimilarity). A: duodenum, B: jejunum, C: ileum and D: caeca.

It is clear that time is a large factor affecting the intestinal microbiota composition, with the day 1 timepoint clustering particularly separately from other timepoints (**Figure 2**). It is also clear that position in the digestive tract is important, with caecal samples clearly separating from other all other parts of the digestive tract throughout development, except at day 14 (**Figure 1**).

At day 1 in jejunal, ileal and caecal samples the most abundant genus on average was *Clostridium_sensu_stricto_1* (jejunum: 0.76 ± 0.32, ileum: 0.86 ± 0.22, caeca 0.95 ± 0.13). The single duodenum sample was dominated by *Enterococcus* (0.86) with smaller amounts of *Clostridium_sensu_stricto_1* (0.14) (**Figure 3**). Sample types did cluster significantly separately from one another by their microbiota compositions (Permutational multivariate analysis of variance (PERMANOVA): P= 0.00478) (**Figure 1**); however, it is not possible to work out which groups of samples may be driving this clustering as after performing pairwise-PERMANOVAs, no sample type clustered significantly separately from another sample type by their community compositions (P > 0.05). However, microbial communities from jejunal and caecal samples were found to be significantly differently rich (P= 0.0018) and diverse (P = 0.033), with jejunum samples having greater diversity and richness (**Figures 4 and 5**). Several OTUs were also found to be more abundant in the small intestinal samples than in caecal samples (**Additional file 4**).

**Figure 3:**
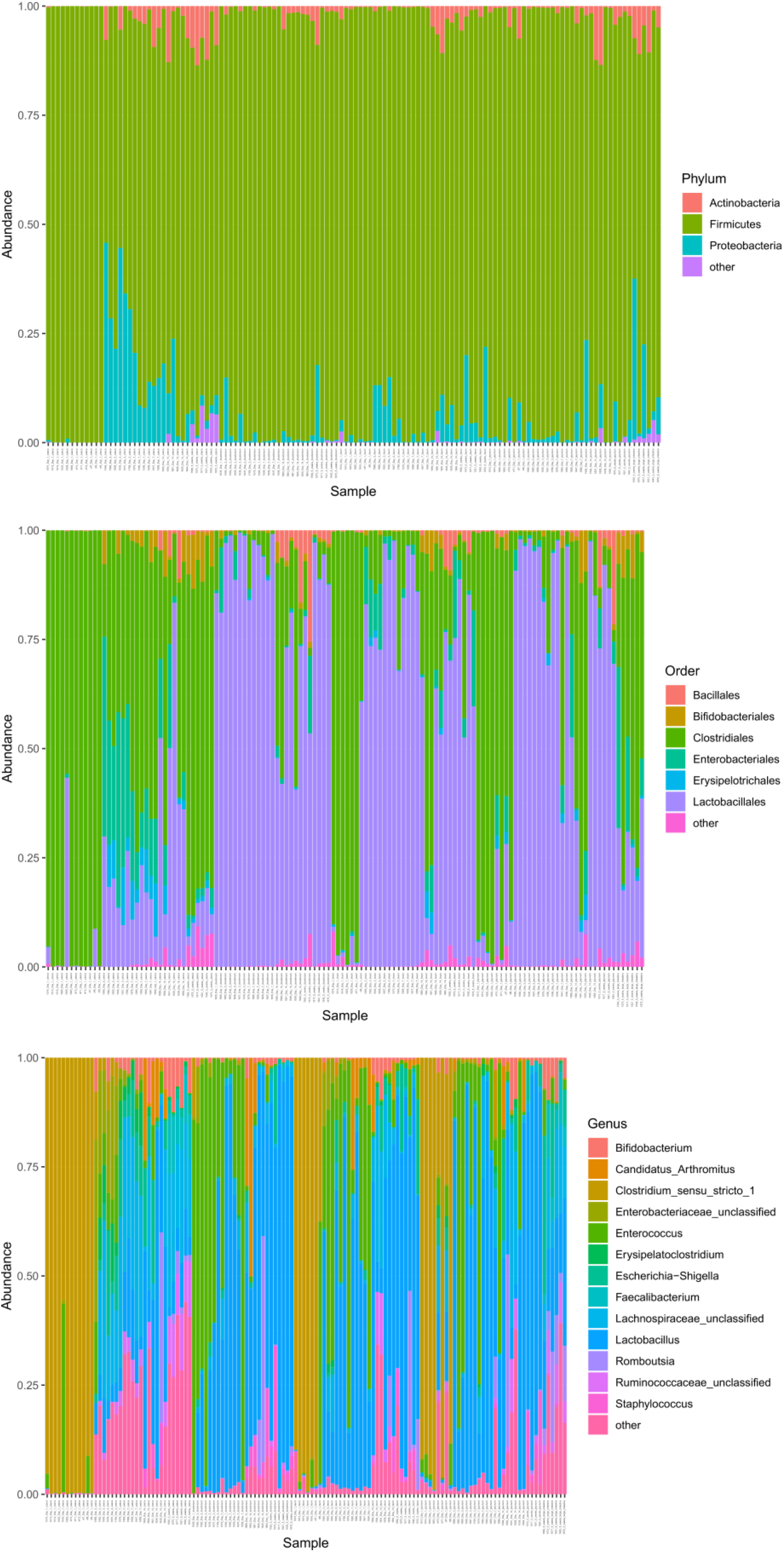
Abundance of bacterial phyla, orders and genera within chicken intestinal samples (subsampled to 10,000 reads). A) Duodenum B) Jejunum C) Ileum D) Caeca.

**Figure 4:**
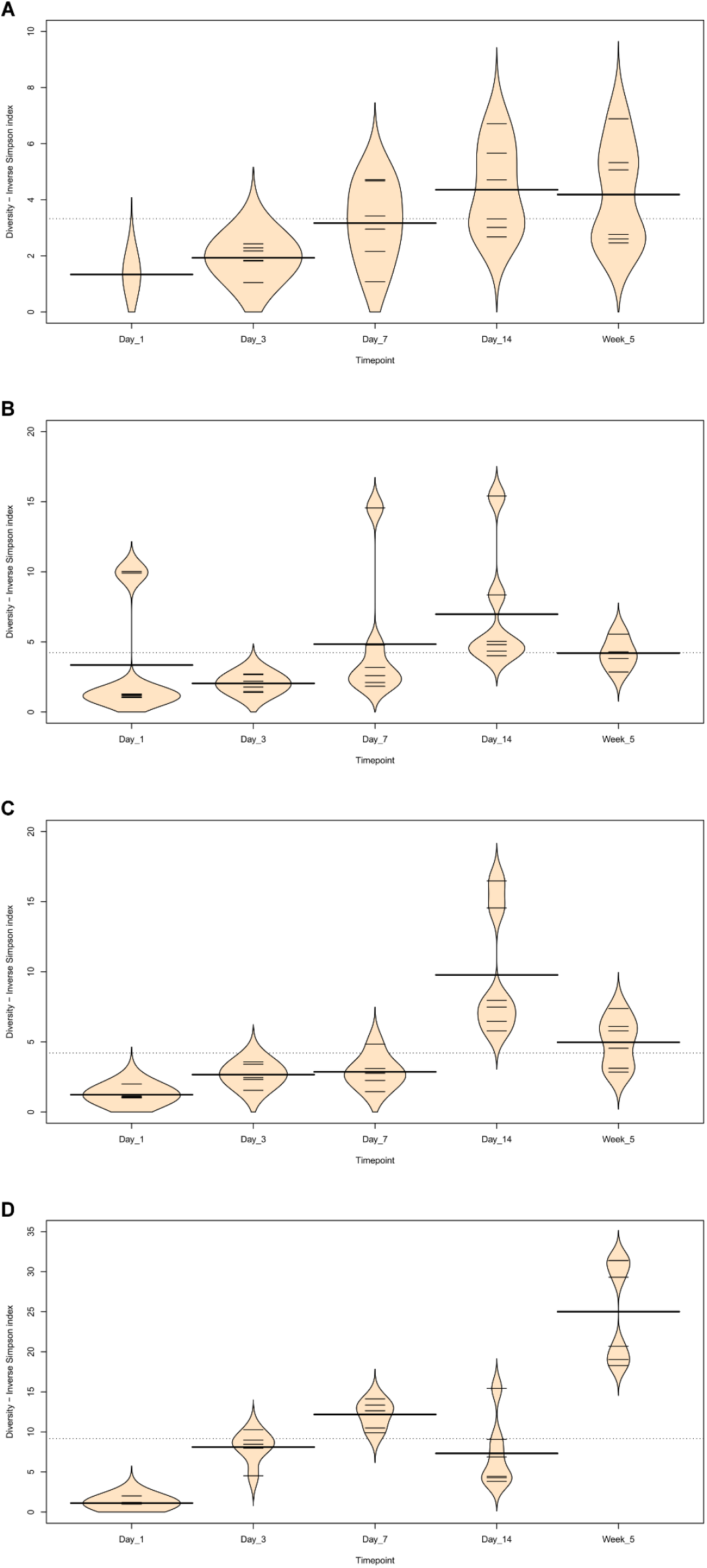
Beanplots showing diversity (Inverse Simpson Index) of chicken intestinal microbiota communities across timepoints (subsampled to 10,000 reads). A) Duodenum B) Jejunum C) Ileum D) Caeca.

**Figure 5:**
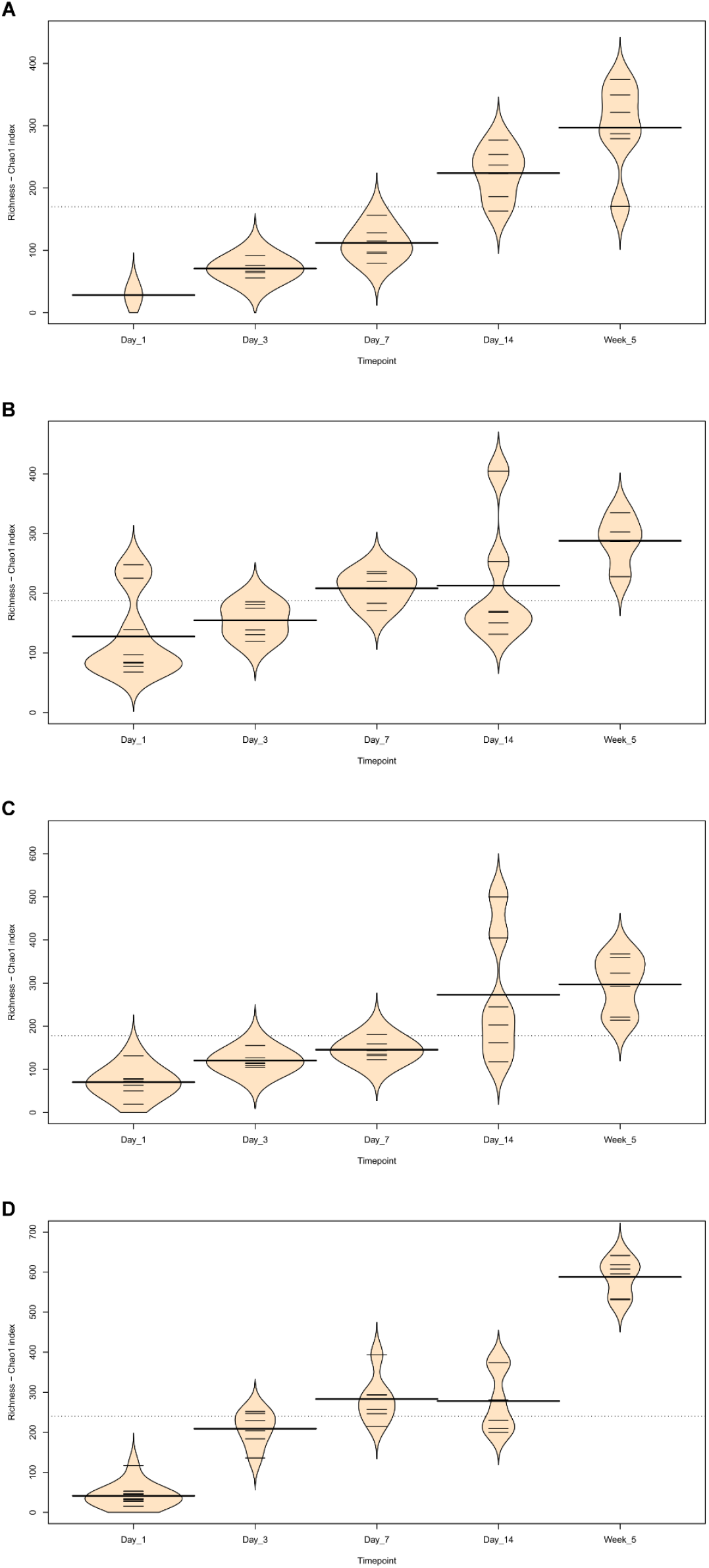
Beanplots showing richness (Chao1 Index) of chicken intestinal microbiota communities across timepoints (subsampled to 10,000 reads). A) Duodenum B) Jejunum C) Ileum D) Caeca.

Between day 1 and day 3, a significant difference was observed in the microbiota composition of all sample types (jejunum: P = 0.01, ileum: P = 0.03, caeca: P = 0.01) except duodenal, probably due to the lack of duodenal samples at the day 1 timepoint. In caecal samples this correlates with a decrease in the abundance of several OTUs including Otu0001 (*Clostridium_sensu_stricto_1*) and an increase in several OTUs belonging to the order *Clostridiales* (**Additional file 5**). This significant decrease in Otu0001 was also observed in duodenal, jejunal and ileal samples (**Additional files 6-8**). Significant increases were found in caecal microbiota richness and ileal and caecal microbiota diversity between these timepoints (Richness - caeca: P = 0.0011; Diversity - ileum: P = 0.023 caeca: P = 0.0011) but not for duodenal or jejunal samples.

At day 3, caecal samples clustered significantly separately by PERMANOVA from all other sample types (duodenum: P = 0.018, jejunum: P = 0.006, ileum: P = 0.024) (**Figure 1**). Duodenal, jejunal and ileal samples did not cluster significantly separately; however, a small number of OTUs were found to be differently abundant (**Additional file 9**). The most common genera found in caecal samples were *Enterobacteriaceae*_unclassified (0.24 ± 0.010), *Lachnospiraceae*_unclassified (0.19 ± 0.13), *Enterococcus* (0.11 ± 0.058), *Escherichia-Shigella* (0.080 ± 0.087) and *Lactobacillus* (0.064 ± 0.033). The most common genera found in ileal samples were *Lactobacillus* (0.47 ± 0.29), *Enterococcus* (0.34 ± 0.26) and *Enterobacteriaceae*_unclassified (0.063 ± 0.045). Duodenal and jejunal samples were both dominated by two genera: *Enterococcus* (duodenum: 0.63 ± 0.23, jejunum: 0.34 ± 0.35) and *Lactobacillus* (duodenum: 0.30 ± 0.24, jejunum: 0.61 ± 0.34). Caecal samples at day 3 were significantly more diverse than other sample types (duodenum: P = 0.013, jejunum: P = 0.013, ileum: P = 0.013) but were only significantly more rich than ileal (P = 0.026) and duodenal (P = 0.013) samples. There was a significant difference in richness but not diversity between duodenal samples and jejunal (richness: P = 0.013) and ileal samples (richness: P = 0.013); no significant differences in richness or diversity were observed between jejunal and ileal samples.

Between day 3 and day 7, no significant differences could be observed in the microbiota composition (PERMANOVA), diversity or richness of any sample type except for an increase in richness in duodenum samples (P = 0.043) and an increase in diversity in caecal samples (P = 0.0433). However, some significant changes in the abundance of specific OTUs were observed. 37 OTUs were more abundant in day 7 caecal samples, the vast majority of which were members of the *Clostridiales* (**Additional file 5**). There was also a significant increase in several members of the *Clostridiales* in duodenal, jejunal and ileal samples along with several Lactobacilli (**Additional files 6-8**).

At day 7, caecal samples still clustered significantly separately by PERMANOVA from all other sample types (duodenum: P = 0.024, jejunum: P = 0.018, ileum: P = 0.024) (**Figure 1**), reflected in the large amount of OTUs which were found to be significantly differently abundant between caecal samples and other sample types (**Additional file 10**). Duodenal, jejunal and ileal samples still did not cluster significantly separately from each other by PERMANOVA. The most common genera found in caecal samples were *Lachnospiraceae*_unclassified (0.34 ± 0.031), *Lactobacillus* (0.117 ± 0.073), *Escherichia-Shigella* (0.10 ± 0.052), *Lachnoclostridium* (0.071 ± 0.031) and *Erysipelatoclostridium* (0.063 ± 0.035). Duodenal, jejunal and ileal samples were all dominated by *Lactobacillus* (duodenum: 0.57 ± 0.35, jejunum: 0.5 ± 0.33, ileum: 0.46 ± 0.26) and *Enterococcus* (duodenum: 0.36 ± 0.35, jejunum: 0.28 ± 0.33, ileum: 0.41 ± 0.32). Caecal samples remained significantly more diverse than duodenal (P = 0.013) and ileal (P = 0.013) samples, but not jejunal samples. There remained a significant difference in richness but not diversity between duodenal samples and jejunal (richness: P = 0.013) samples but not ileal samples. A significant difference in the richness but not diversity of samples was observed between the jejunum and ileum (richness: P = 0.026).

Between day 7 and day 14, significant changes in community composition were observed for caecal (P = 0.03), ileal (P = 0.05) and duodenal (P = 0.03) samples; a difference in diversity was observed in ileal samples (P = 0.022); and differences in richness were observed in duodenal (P = 0.022) and jejunal (P = 0.022) samples. Several OTUs were also found to have changed in abundance within sample types (**Additional file 5-8**). These changes appear to have led to increased homogeneity of microbial communities across all sample types (**Figure 1**). No significant differences in community composition, richness or diversity were observed between the sample types at this timepoint and very few specific OTUs were found to be different between sample types (**Additional file 11**). This seems to have been driven by caecal samples attaining a “small-intestine like” microbiota dominated by *Lactobacillus* (0.39 ± 0.24) with smaller numbers of *Lachnospiraceae*_unclassified (0.10 ± 0.13), *Candidatus_Arthromitus* (0.1 ± 0.087), *Escherichia-Shigella* (0.098 ± 0.093) and *Romboutsia* (0.073 ± 0.018). The most common genera in dudodenal samples were *Lactobacillus* (0.54 ± 0.17), *Candidatus_Arthromitus* (0.15 ± 0.20), *Romboutsia* (0.10 ± 0.21) and *Staphylococcus* (0.056 ± 0.053); in jejunal samples were *Lactobacillus* (0.38 ± 0.31), *Lachnospiraceae*_unclassified (0.16 ± 0.19), *Romboutsia* (0.073 ± 0.18), *Enterococcus* (0.072 ± 0.15) and *Escherichia-Shigella* (0.058 ±0.094); and in ileal samples were *Lactobacillus* (0.42 ± 0.28), *Lachnospiraceae*_unclassified (0.17 ± 0.12), *Ruminococcaceae*_unclassified (0.057 ± 0.058).

Between day 14 and 5 weeks, a significant difference can be observed in the microbiota composition of the ileum (P = 0.04, **Additional file 8**) and caeca (P = 0.03, **Additional file 5**) but not for jejunal and duodenal samples, despite significant changes in the abundance of specific OTUs (**Additional files 6 and 7**). Significant increases in richness and diversity were observed for caecal samples (P = 0.0216). The most common genera in caecal samples were *Lachnospiraceae*_unclassified (0.23 ± 0.050), *Ruminococcaceae*_unclassified (0.11 ± 0.016), *Faecalibacterium* (0.11 ± 0.022), *Bifidobacterium* (0.080 ± 0.041), *Lactobacillus* (0.066 ± 0.034), *Clostridiales_vadinBB60_group_ge* (0.055 ± 0.027). Ileal, duodenal and jejunal samples were predominated by *Lactobacillus* (duodenum: 0.81 ± 0.19, jejunum: 0.77 ± 0.11, ileum: 0.65 ± 0.20) with the next most common genus being *Romboutsia* (0.090 ± 0.15) in ileal samples and *Staphylococcus* in duodenal (0.060 ± 0.090) and jejunal (0.097 ± 0.090) samples. Large intestine samples included genera found in both small intestinal and caecal samples: *Lactobacillus* (0.21 ± 0.074), *Lachnospiraceae*_unclassified (0.15 ± 0.066), *Escherichia-Shigella* (0.12 ± 0.14), *Romboutsia* (0.092 ± 0.071), *Faecalibacterium* (0.067 ± 0.043), *Ruminococcaceae*_unclassified (0.061 ± 0.036) and *Bifidobacterium* (0.060 ± 0.036).

A significant difference was found by PERMANOVA between caecal samples and duodenal (P = 0.04) and ileal (P = 0.02) samples but not large intestine or jejunal samples (**Figure 1**). The bacterial community composition of large intestine samples was found to be significantly different to all other sample types except the caeca (duodenum: P = 0.03, jejunum: P = 0.04, ileum: P = 0.04). This is reflected in our Deseq analysis where no OTUs were found to be significantly differently abundant between large intestine and caecal samples but many OTUs were differently abundant between the large intestine and other sample types (**Additional file 12**). No significant differences by PERMANOVA were observed between duodenal, jejunal and ileal samples, although several OTUs were found to be significantly differently abundant (**Additional file 12**). Diversity and richness were significantly higher in caecal samples than duodenal (P = 0.022) or ileal (P = 0.022) but not jejunal samples, although this may be due to a lack of jejunal samples at this timepoint as two of the six samples had to be discarded due to a lack of reads.

## Discussion

In this study we compared samples taken from the duodenum, jejunum, ileum and caeca of broiler chickens at five timepoints from day of hatch to 5 weeks of age. Gaining an insight into how the microbiota changes over time at specific sites in the small intestine may lead us to a better understanding of the microbial ecology of the chicken gut. We found changes in community composition, diversity and richness for all sample types over time. More specifically, we observed an increase in the richness of microbial communities in all gut sections, and a general increase in diversity except at the day 14 timepoint. While the succession of communities was different in each section of the gut, we can broadly say that the intestinal microbiota of our chickens was initially formed by a low diversity community of predominantly *Clostridium_sensu_stricto_1* which diversified over time to contain a far greater variety of *Clostridiales*, with smaller numbers of other taxonomies. Samples taken from different locations within the small intestine were found to be similar to one another, whereas in the majority of timepoints caecal samples clustered significantly separately by composition from small intestinal samples. However, at several timepoints specific OTUs were found to be significantly more abundant in certain small intestinal locations in comparison to other sites, suggesting that there are differences between the microbial communities across the small intestine and therefore a sample taken from one site should not be taken as representative of the small intestine as a whole.

At the day 1 timepoint samples from the caeca and the small intestine were dominated by *Clostridium_sensu_stricto_1*. The lack of diversity observed at this timepoint was expected as previous studies have also shown that microbial communities in the chicken intestine in the first days of life have little diversity and are usually dominated by either members of the *Enterobacteriaceae* or the *Clostridiaceae* [9–11, 14, 15, 26]. It is possible that members of these communities act as founding species for chicken gut microbial communities. It has been demonstrated that these early life communities can have a significant impact on later microbiota communities and bird phenotypes [10, 23] and it is therefore important to understand the origin of these founder microbes and the impact they may have.

Between day 1 and day 3 the proportion of *Clostridium_sensu_stricto_1* greatly decreased in all of our sample types and while it remained present at later timepoints it was never highly abundant. At day 3 all small intestinal sample types were dominated by two genera: *Lactobacillus* and *Enterococcus*. These would remain the dominant genera in small intestinal samples at all future timepoints. *Lactobacillus* has previously been noted as being highly abundant in the small intestine [12, 14, 17, 18, 20, 23] as has *Enterococcus* [14]. In our study at this timepoint the caeca were dominated by *Enterococcus*, *Escherichia-Shigella*, *Lactobacillus* and unclassified members of the *Enterobacteriaceae* and the *Lachnospiraceae*.

No significant change was observed in the community composition of any sample type between day 3 and 7; however, at timepoint 14 days changes were observed in the caecal microbial communities which led them to no longer cluster separately from small intestinal samples by their community compositions. We are unsure of the cause of this seeming “homogenisation” of our samples at this timepoint and this phenomenon does not previously appear in the chicken literature. This homogenisation correlated with an increase in the relative abundance of members of *Lactobacillus* and *Candidatus Arthromitus* which have previously been associated with the small intestine [12, 23]. By the 5 week timepoint further changes in the microbial community composition led to the caecal microbiota again clustering separately from all other sample types.

We ran several reagent only controls alongside our samples. The most abundant genera in these controls were also found in our samples: *Clostridium_sensu_stricto_1*, *Lactobacillus*, *Enterococcus* and unclassified members of the *Lachnospiraceae*. It seems unlikely that the presence of these bacteria is due to reagent contamination as they are common intestinal colonisers, unlike the skin and environmental bacteria which tend to occur as contaminants in reagents [27]. Two possible alternative explanations for the presence of these bacteria are index switching or cross-well contamination [28, 29]. Due to the low numbers of reads seen in the negative controls, contamination is unlikely to have driven any of the observed differences, though as always with microbiota studies, our results should be interpreted with care.

## Conclusions

While there is high variability present between the microbiota of chickens in different trials [21] it does seem as though there are taxonomies which are consistently present in the chicken gut [1]. Identifying these shared taxa and elucidating their function within birds could lead to a greater understanding of the microbial ecology of the chicken gut and the development of technologies which could be widely applied to improve chicken health or productivity.

## Methods

### Ethics approval

Animals were housed in premises licensed under a UK Home Office Establishment License within the terms of the UK Home Office Animals (Scientific Procedures) Act 1986. Housing and husbandry complied with the Code of Practice for Housing and Care of Animals Bred, Supplied or Used for Scientific Purposes and were overseen by the Roslin Institute Animal Welfare and Ethical Review Board. Animals were culled by schedule one methods authorized by the Animals (Scientific Procedures) Act 1986. Birds were euthanized by cervical dislocation.

### Study design

Ross 308 (Aviagen) chickens were hatched and housed at the National Avian Research Facility in Edinburgh (UK). Chickens were housed in groups in floor pens with wood shaving bedding and received food and water ad libitum. Chickens received the Marek’s-Rispens vaccine and feed included coccidiostats. Samples were collected from chickens at the following ages: day 1 (n=12), day 3 (n=6), day 7 (n=6), day 14 (n=6) and day 35 (5 weeks: n=6)). Collected samples included intestinal contents from the duodenum, jejunum, ileum and caeca. Large intestinal contents were also collected at 5 weeks.

Our experiments followed the general principles set out in Pollock *et al.* [30]. Samples were stored at 4°C for a maximum of 24 hours until DNA extraction, except for those from DNA extraction batch 11 which were frozen at −20°C for 9 days prior to DNA extraction. The extraction batches to which each sample belonged can be found in **Additional file 13**. DNA extraction was performed as described previously using the DNeasy PowerLyzer PowerSoil Kit (Qiagen) [31]. Reagent only controls were included for each batch of DNA extractions. DNA was also extracted from a mock community control (20 Strain Even Mix Whole Cell Material (ATCC^®^ MSA-2002™)). The V4 region of the 16S rRNA gene was amplified from the extracted DNA, following a previously described method [32]. The sequencing reaction consisted of 12.5 μl of Q5 High Fidelity DNA Polymerase (New England Biolabs), 1.25 μl each of custom 10nM barcoded forward (5’–TATGGTAATTGTGTGCCAGCMGCCGCGGTAA–3’) and reverse primers (5’–AGTCAGTCAGCCGGACTACHVGGGTWTCTAAT–3’), 9 μl of nuclease free water (Qiagen) and 1 μl of DNA template. Cycling conditions were: 95°C for 2 min followed by 30 cycles of 95°C for 20 sec, 55°C for 15 sec, 72°C for 5 min followed by 72°C for 10 min. Amplicons were purified using the AMPure XP System (Beckman Coulter), according to the manufacturer’s instructions, except that a 1:1 ratio of AMPure beads to sample was used. The amplicon concentrations were determined using the Qubit 3.0 Fluorometer (Life Technologies) with the Qubit dsDNA HS Assay Kit (Life Technologies), according to the manufacturer’s instructions. Samples were then pooled into an equimolar library, which was sequenced on an Illumina MiSeq v.2 (Illumina Inc.) producing 250 bp reads ( 1^st^ read primer: 5’–TATGGTAATTGTGTGCCAGCMGCCGCGGTAA–3’, 2^nd^ read primer: 5’– AGTCAGTCAGCCGGACTACHVGGGTWTCTAAT–3’, index primer: 5’– ATTAGAWACCCBDGTAGTCCGGCTGACTGACT–3’.

### Bioinformatics

Mothur [33] was used for quality control of sequences, alignment, taxonomic assignment and OTU clustering, following a modified version of the MiSeq pipeline supplied on the mothur website [32]. Sequences were removed if they were >275 bp in length, contained ambiguous bases, had homopolymers of >9bp in length, did not align to the V4 region of the 16S rRNA gene or they did not originate from bacteria. Chimeras were identified using UCHIME [34] and were removed. The SILVA database [35], trimmed to the V4 16S rRNA gene, was used for alignment and taxonomic identification of sequences. Where the taxonomy of an OTU is labelled as ###_unclassified, this indicates that the OTU was unable to be identified to that level of taxonomy and the ### indicates the lowest taxonomy this OTU could be assigned to. OTUs were clustered by similarity using the dist.seqs and cluster commands from within mothur, using the default parameters.

Richness and diversity were calculated within mothur. Community richness was measured using the Chao 1 index. Community diversity was measured using the Inverse Simpson’s Index. The following analyses were performed in R (version 3.5.1.) [36] with the seed 8765. To compare the diversity and richness of microbial communities between groups, the Pairwise Wilcoxon Rank Sum Test was used with Bonferroni corrections to correct for multiple comparisons. Non-metric Multidimensional Scaling (NMDS) graphs were constructed using the Vegan [37] package and ggplot2 [38], using the Bray–Curtis dissimilarity. UpSet graphs were constructed using the UpSetR package [39]. Beanplots were constructed using the beanplot package [40]. PERMANOVA analyses were performed using the adonis function from the Vegan package; pairwise comparisons were performed using an adjusted version of the adonis function which output pairwise comparisons with Bonferroni corrections to correct for multiple comparisons. The package DESeq2 [41] was used to calculate differences in abundances between groups for individual OTUs. Prior to statistical analysis, samples were subsampled to 10,000 reads, except for DEseq2 analysis.

### Availability of data and materials

The paired-read fastq data that support the findings of this study have been deposited in in the European Nucleotide Archive with the accession code PRJEB33615.

## Acknowledgements

We would like to thank the staff at the Greenwood Building, Roslin Institute for the care of our animals. Sequencing was carried out by Edinburgh Genomics, The University of Edinburgh. The Roslin Institute forms part of the Royal (Dick) School of Veterinary Studies, University of Edinburgh. This project was supported by the Biotechnology and Biological Sciences Research Council, including institute strategic programme and national capability awards to The Roslin Institute (BBSRC: BB/P013759/1, BB/P013732/1, BB/J004235/1, BB/J004243/1). The funding body had no role in the design of the study and collection, analysis, and interpretation of data and in writing the manuscript.

## Additional material

**Additional file 1** (Additional file 1.xlsx): Abundance of OTUs in chicken intestinal samples (not sub-sampled).

**Additional file 2** (Additional file 2.xlsx): Taxonomy of bacterial OTUs.

**Additional file 3** (Additional file 3.docx): Contains Table 1 (Mock community composition by genus), Table 2 (Mock community composition by order), Table 3 (Diversity (Inverse Simpson) and richness (Chao 1 index) of subsampled samples), Figure 1 (UpSet graph – timepoints) and Figure 1 (UpSet graph – timepoints).

**Additional file 4** (Additional file 4.xlsx): OTUs which were identified as being significantly more abundant by DESeq2 between locations within the chicken intestinal tract at day 1 of age.

**Additional file 5** (Additional file 5.xlsx): OTUs which were identified as being significantly more abundant by DESeq2 between chicken caecal samples at different timepoints.

**Additional file 6** (Additional file 6.xlsx): OTUs which were identified as being significantly more abundant by DESeq2 between chicken duodenal samples at different timepoints.

**Additional file 7** (Additional file 7.xlsx): OTUs which were identified as being significantly more abundant by DESeq2 between chicken jejunal samples at different timepoints.

**Additional file 8** (Additional file 8.xlsx): OTUs which were identified as being significantly more abundant by DESeq2 between chicken ileal samples at different timepoints.

**Additional file 9** (Additional file 9.xlsx): OTUs which were identified as being significantly more abundant by DESeq2 between locations within the chicken intestinal tract at day 3 of age.

**Additional file 10** (Additional file 10.xlsx): OTUs which were identified as being significantly more abundant by DESeq2 between locations within the chicken intestinal tract at day 7 of age.

**Additional file 11** (Additional file 11.xlsx): OTUs which were identified as being significantly more abundant by DESeq2 between locations within the chicken intestinal tract at day 14 of age.

**Additional file 12** (Additional file 12.xlsx): OTUs which were identified as being significantly more abundant by DESeq2 between locations within the chicken intestinal tract at 5 weeks of age.

**Additional file 13** (Additional file 13.xlsx): Sample metadata.

