## Additional file 3 for "Development of the duodenal, ileal, jejunal and caecal microbiota in chickens"

**Table 1: Mock community composition by genus**

| **Genus ID** | **Expected abundance** | **Actual abundance** |
| --- | --- | --- |
| *Acinetobacter* | 0.05 | 0.12 |
| *Actinomyces* | 0.05 | 0.0042 |
| *Bacillus* | 0.05 | 0 |
| *Bacteroides* | 0.05 | 0.039 |
| *Bifidobacterium* | 0.05 | 0.0046 |
| *Clostridium* | 0.05 | 0.022 |
| *Cutibacterium* | 0.05 | 0.00033 |
| *Deinococcus* | 0.05 | 0.033 |
| *Enterococcus* | 0.05 | 0.046 |
| *Escherichia* | 0.05 | 0.075 |
| *Helicobacter* | 0.05 | 0.00086 |
| *Lactobacillus* | 0.05 | 0.032 |
| *Neisseria* | 0.05 | 0.059 |
| *Porphyromonas* | 0.05 | 0.016 |
| *Pseudomonas* | 0.05 | 0.07 |
| *Rhodobacter* | 0.05 | 0.018 (as family *Rhodobacteraceae*_unclassified) |
| *Staphylococcus* | 0.15 | 0.054 |
| *Streptococcus* | 0.1 | 0.2 |

**Table 2: Mock community composition by order**

| **Taxonomy** | **Expected abundance** | **Actual abundance** |
| --- | --- | --- |
| *Actinomycetales* | 0.1 | 0.0042 |
| *Bacillales* | 0.2 | 0.24 |
| *Bacteroidales* | 0.1 | 0.055 |
| *Bifidobacteriales* | 0.05 | 0.0046 |
| *Campylobacterales* | 0.05 | 0.00086 |
| *Clostridiales* | 0.05 | 0.031 |
| *Deinococcales* | 0.05 | 0.033 |
| *Enterobacteriales* | 0.05 | 0.075 |
| *Lactobacillales* | 0.2 | 0.28 |
| *Betaproteobacteriales* | 0.05 | 0.059 |
| *Pseudomonadales* | 0.1 | 0.19 |
| *Rhodobacterales* | 0.05 | 0.018 |

**Table 3: Diversity (Inverse Simpson) and richness (Chao 1 index) of subsampled samples**

| **Sample** | **Inverse Simpsons** | **Inverse Simpsons (lower confidence interval)** | **Inverse Simpsons (higher confidence interval)** | **Chao 1 index** | **Chao 1 index (lower confidence interval)** | **Chao 1 index (higher confidence interval)** |
| --- | --- | --- | --- | --- | --- | --- |
| 1574_Day_1_caeca | 1.11 | 1.10 | 1.12 | 116.66 | 80.07 | 210.52 |
| 1613_Day_1_caeca | 1.00 | 1.00 | 1.00 | 28.13 | 15.55 | 79.94 |
| 1614_Day_1_caeca | 1.01 | 1.00 | 1.01 | 45.26 | 27.02 | 110.71 |
| 1620_Day_1_caeca | 1.00 | 1.00 | 1.00 | 15.45 | 8.66 | 50.08 |
| 1635_Day_1_caeca | 2.02 | 2.01 | 2.03 | 53.34 | 29.11 | 139.64 |
| 1663_Day_1_caeca | 1.00 | 1.00 | 1.00 | 32.31 | 16.48 | 93.98 |
| nf10_Day_1_caeca | 1.01 | 1.01 | 1.01 | 27.40 | 17.11 | 71.43 |
| nf11_Day_1_caeca | 1.00 | 1.00 | 1.01 | 47.01 | 25.52 | 123.64 |
| nf12_Day_1_caeca | 1.01 | 1.00 | 1.01 | 52.32 | 29.00 | 133.13 |
| nf7_Day_1_caeca | 1.00 | 1.00 | 1.01 | 33.93 | 18.55 | 94.33 |
| nf8_Day_1_caeca | 1.19 | 1.18 | 1.21 | 15.37 | 10.16 | 44.07 |
| nf9_Day_1_caeca | 1.01 | 1.00 | 1.01 | 29.25 | 19.65 | 69.54 |
| 1635_Day_1_duodenum | 1.34 | 1.32 | 1.36 | 28.15 | 19.50 | 66.77 |
| 1574_Day_1_ileum | 1.25 | 1.23 | 1.26 | 131.50 | 114.61 | 170.77 |
| 1613_Day_1_ileum | 1.07 | 1.06 | 1.07 | 50.29 | 43.13 | 75.49 |
| 1614_Day_1_ileum | 1.11 | 1.10 | 1.12 | 76.90 | 67.14 | 106.76 |
| 1620_Day_1_ileum | 1.01 | 1.01 | 1.02 | 19.42 | 17.66 | 30.78 |
| 1635_Day_1_ileum | 1.18 | 1.17 | 1.19 | 62.81 | 50.57 | 102.15 |
| nf11_Day_1_ileum | 1.04 | 1.04 | 1.05 | 78.59 | 57.93 | 138.63 |
| nf8_Day_1_ileum | 2.00 | 1.98 | 2.02 | 71.98 | 57.23 | 114.86 |
| 1613_Day_1_jejunum | 1.17 | 1.16 | 1.19 | 96.64 | 90.71 | 114.62 |
| 1614_Day_1_jejunum | 1.22 | 1.21 | 1.24 | 77.29 | 71.17 | 100.00 |
| 1620_Day_1_jejunum | 1.10 | 1.09 | 1.10 | 67.73 | 60.58 | 91.68 |
| 1663_Day_1_jejunum | 1.02 | 1.02 | 1.03 | 139.07 | 83.17 | 286.84 |
| nf10_Day_1_jejunum | 10.01 | 9.67 | 10.38 | 247.96 | 228.23 | 287.57 |
| nf11_Day_1_jejunum | 1.08 | 1.08 | 1.09 | 82.94 | 66.66 | 127.21 |
| nf7_Day_1_jejunum | 9.91 | 9.55 | 10.30 | 225.10 | 206.81 | 264.04 |
| nf8_Day_1_jejunum | 1.28 | 1.26 | 1.30 | 84.27 | 70.16 | 120.97 |
| 1569_Day_14_caeca | 4.27 | 4.18 | 4.36 | 209.33 | 158.23 | 311.42 |
| 1594_Day_14_caeca | 15.43 | 14.93 | 15.97 | 373.81 | 292.83 | 523.78 |
| 1601_Day_14_caeca | 6.89 | 6.75 | 7.03 | 229.81 | 174.56 | 339.50 |
| 1626_Day_14_caeca | 3.83 | 3.74 | 3.94 | 199.78 | 148.76 | 304.88 |
| 1639_Day_14_caeca | 4.41 | 4.27 | 4.56 | 373.75 | 305.21 | 488.88 |
| 1664_Day_14_caeca | 9.08 | 8.78 | 9.40 | 280.41 | 214.15 | 414.98 |
| 1569_Day_14_duodenum | 5.68 | 5.53 | 5.83 | 222.78 | 172.24 | 324.60 |
| 1594_Day_14_duodenum | 3.06 | 2.99 | 3.13 | 188.83 | 137.51 | 297.51 |
| 1601_Day_14_duodenum | 4.74 | 4.60 | 4.90 | 278.72 | 221.43 | 385.63 |
| 1626_Day_14_duodenum | 2.69 | 2.62 | 2.76 | 237.58 | 187.70 | 333.04 |
| 1639_Day_14_duodenum | 3.30 | 3.21 | 3.39 | 162.33 | 135.75 | 219.47 |
| 1664_Day_14_duodenum | 6.68 | 6.53 | 6.84 | 253.74 | 206.10 | 342.26 |
| 1569_Day_14_ileum | 7.44 | 7.27 | 7.63 | 244.89 | 195.37 | 343.88 |
| 1594_Day_14_ileum | 14.57 | 13.93 | 15.27 | 412.38 | 318.95 | 581.33 |
| 1601_Day_14_ileum | 16.52 | 16.02 | 17.06 | 504.95 | 378.71 | 727.83 |
| 1626_Day_14_ileum | 7.99 | 7.77 | 8.21 | 156.90 | 137.34 | 203.66 |
| 1639_Day_14_ileum | 6.46 | 6.24 | 6.69 | 205.21 | 170.95 | 278.75 |
| 1664_Day_14_ileum | 5.69 | 5.55 | 5.85 | 113.49 | 89.92 | 192.82 |
| 1569_Day_14_jejunum | 5.03 | 4.91 | 5.15 | 169.43 | 137.63 | 239.39 |
| 1594_Day_14_jejunum | 4.32 | 4.19 | 4.47 | 253.34 | 221.33 | 314.36 |
| 1601_Day_14_jejunum | 8.34 | 8.02 | 8.69 | 150.23 | 124.33 | 220.00 |
| 1626_Day_14_jejunum | 15.36 | 14.84 | 15.93 | 404.66 | 315.37 | 567.18 |
| 1639_Day_14_jejunum | 3.99 | 3.92 | 4.07 | 131.13 | 97.51 | 212.19 |
| 1664_Day_14_jejunum | 4.80 | 4.73 | 4.87 | 167.92 | 131.71 | 244.74 |
| 1566_Day_3_caeca | 4.52 | 4.39 | 4.66 | 136.66 | 102.66 | 214.73 |
| 1582_Day_3_caeca | 8.46 | 8.24 | 8.70 | 204.23 | 134.53 | 372.56 |
| 1605_Day_3_caeca | 8.98 | 8.79 | 9.19 | 184.54 | 134.11 | 299.23 |
| 1636_Day_3_caeca | 8.47 | 8.23 | 8.72 | 247.57 | 175.05 | 403.12 |
| 1637_Day_3_caeca | 7.97 | 7.72 | 8.23 | 229.44 | 173.31 | 344.98 |
| 1769_Day_3_caeca | 10.27 | 9.98 | 10.57 | 251.56 | 183.06 | 394.52 |
| 1566_Day_3_duodenum | 2.28 | 2.23 | 2.34 | 70.93 | 54.15 | 124.65 |
| 1582_Day_3_duodenum | 1.82 | 1.79 | 1.85 | 56.06 | 40.92 | 111.26 |
| 1605_Day_3_duodenum | 1.05 | 1.04 | 1.05 | 75.61 | 58.75 | 122.90 |
| 1636_Day_3_duodenum | 2.43 | 2.37 | 2.49 | 91.55 | 67.83 | 159.52 |
| 1637_Day_3_duodenum | 2.17 | 2.14 | 2.21 | 66.95 | 47.02 | 130.23 |
| 1769_Day_3_duodenum | 1.84 | 1.81 | 1.87 | 63.75 | 43.99 | 130.48 |
| 1566_Day_3_ileum | 3.41 | 3.34 | 3.48 | 104.03 | 74.12 | 187.44 |
| 1582_Day_3_ileum | 2.70 | 2.63 | 2.77 | 155.06 | 109.23 | 267.29 |
| 1605_Day_3_ileum | 2.31 | 2.25 | 2.37 | 109.17 | 83.04 | 175.45 |
| 1636_Day_3_ileum | 3.60 | 3.49 | 3.71 | 126.58 | 93.28 | 214.12 |
| 1637_Day_3_ileum | 2.46 | 2.43 | 2.50 | 112.49 | 87.07 | 176.79 |
| 1769_Day_3_ileum | 1.53 | 1.50 | 1.56 | 114.88 | 84.66 | 192.19 |
| 1566_Day_3_jejunum | 2.67 | 2.62 | 2.72 | 174.13 | 134.82 | 258.65 |
| 1582_Day_3_jejunum | 1.40 | 1.38 | 1.42 | 118.99 | 89.28 | 192.37 |
| 1605_Day_3_jejunum | 1.79 | 1.76 | 1.82 | 138.20 | 102.82 | 223.99 |
| 1636_Day_3_jejunum | 2.19 | 2.15 | 2.24 | 130.67 | 94.34 | 219.16 |
| 1637_Day_3_jejunum | 2.72 | 2.67 | 2.77 | 180.31 | 136.30 | 276.05 |
| 1769_Day_3_jejunum | 1.45 | 1.43 | 1.48 | 185.77 | 137.86 | 289.38 |
| 1579_Day_7_caeca | 9.89 | 9.62 | 10.16 | 214.51 | 155.25 | 346.77 |
| 1585_Day_7_caeca | 13.35 | 12.89 | 13.85 | 393.54 | 305.60 | 550.22 |
| 1599_Day_7_caeca | 12.64 | 12.19 | 13.12 | 294.06 | 228.80 | 419.18 |
| 1609_Day_7_caeca | 12.64 | 12.34 | 12.96 | 256.88 | 184.54 | 410.92 |
| 1647_Day_7_caeca | 14.11 | 13.73 | 14.52 | 292.65 | 216.22 | 444.50 |
| 1658_Day_7_caeca | 10.50 | 10.23 | 10.78 | 246.44 | 189.04 | 362.22 |
| 1579_Day_7_duodenum | 3.42 | 3.32 | 3.53 | 156.53 | 125.45 | 226.57 |
| 1585_Day_7_duodenum | 2.16 | 2.11 | 2.20 | 79.38 | 61.28 | 131.11 |
| 1599_Day_7_duodenum | 2.95 | 2.89 | 3.02 | 128.11 | 102.36 | 188.50 |
| 1609_Day_7_duodenum | 4.68 | 4.62 | 4.74 | 97.27 | 78.24 | 155.49 |
| 1647_Day_7_duodenum | 4.72 | 4.61 | 4.83 | 114.88 | 94.87 | 166.49 |
| 1658_Day_7_duodenum | 1.07 | 1.06 | 1.08 | 94.63 | 62.97 | 184.63 |
| 1579_Day_7_ileum | 1.42 | 1.40 | 1.45 | 122.16 | 92.42 | 197.12 |
| 1585_Day_7_ileum | 4.85 | 4.75 | 4.95 | 181.54 | 137.17 | 275.97 |
| 1599_Day_7_ileum | 2.74 | 2.67 | 2.81 | 132.38 | 101.29 | 207.35 |
| 1609_Day_7_ileum | 2.87 | 2.83 | 2.92 | 142.76 | 109.48 | 218.97 |
| 1647_Day_7_ileum | 3.08 | 3.02 | 3.14 | 135.19 | 105.28 | 202.96 |
| 1658_Day_7_ileum | 2.23 | 2.18 | 2.29 | 158.47 | 117.27 | 254.80 |
| 1579_Day_7_jejunum | 1.86 | 1.82 | 1.91 | 219.52 | 174.25 | 311.55 |
| 1585_Day_7_jejunum | 4.75 | 4.61 | 4.90 | 232.50 | 169.29 | 366.85 |
| 1599_Day_7_jejunum | 2.54 | 2.49 | 2.60 | 207.91 | 157.66 | 314.93 |
| 1609_Day_7_jejunum | 3.16 | 3.09 | 3.24 | 182.41 | 140.44 | 269.76 |
| 1647_Day_7_jejunum | 14.58 | 14.15 | 15.04 | 236.35 | 216.82 | 275.35 |
| 1658_Day_7_jejunum | 2.11 | 2.07 | 2.16 | 171.66 | 128.47 | 270.54 |
| 1562_5_weeks_caeca | 18.28 | 17.61 | 18.99 | 531.15 | 432.00 | 697.55 |
| 1573_5_weeks_caeca | 19.02 | 18.33 | 19.77 | 641.29 | 527.19 | 823.10 |
| 1617_5_weeks_caeca | 31.35 | 30.19 | 32.60 | 607.85 | 512.94 | 759.61 |
| 1621_5_weeks_caeca | 20.69 | 19.93 | 21.52 | 532.63 | 436.83 | 690.16 |
| 1642_5_weeks_caeca | 31.42 | 30.22 | 32.71 | 617.98 | 512.30 | 788.11 |
| 1672_5_weeks_caeca | 29.31 | 28.28 | 30.43 | 595.83 | 488.31 | 771.48 |
| 1562_5_weeks_duodenum | 2.44 | 2.37 | 2.51 | 288.18 | 243.86 | 368.67 |
| 1573_5_weeks_duodenum | 5.32 | 5.20 | 5.44 | 278.24 | 206.76 | 417.29 |
| 1617_5_weeks_duodenum | 5.08 | 5.00 | 5.16 | 168.82 | 127.10 | 258.27 |
| 1621_5_weeks_duodenum | 2.63 | 2.56 | 2.70 | 350.61 | 294.34 | 447.17 |
| 1642_5_weeks_duodenum | 2.76 | 2.70 | 2.81 | 320.15 | 262.57 | 419.16 |
| 1672_5_weeks_duodenum | 6.91 | 6.76 | 7.06 | 374.27 | 321.98 | 461.77 |
| 1562_5_weeks_large_intestine | 6.06 | 5.86 | 6.28 | 309.57 | 256.05 | 407.56 |
| 1573_5_weeks_large_intestine | 20.45 | 19.73 | 21.21 | 637.51 | 526.39 | 813.98 |
| 1617_5_weeks_large_intestine | 9.01 | 8.73 | 9.31 | 392.71 | 328.73 | 503.16 |
| 1621_5_weeks_large_intestine | 17.51 | 16.98 | 18.07 | 526.58 | 421.84 | 701.45 |
| 1642_5_weeks_large_intestine | 26.28 | 25.29 | 27.34 | 582.78 | 490.18 | 731.07 |
| 1672_5_weeks_large_intestine | 16.06 | 15.61 | 16.54 | 484.33 | 396.22 | 632.38 |
| 1562_5_weeks_ileum | 5.82 | 5.67 | 5.98 | 360.46 | 301.18 | 460.78 |
| 1573_5_weeks_ileum | 3.12 | 3.04 | 3.20 | 222.01 | 166.03 | 334.61 |
| 1617_5_weeks_ileum | 6.13 | 6.02 | 6.24 | 294.18 | 245.40 | 380.25 |
| 1621_5_weeks_ileum | 4.59 | 4.48 | 4.71 | 321.89 | 256.85 | 436.89 |
| 1642_5_weeks_ileum | 2.83 | 2.75 | 2.91 | 366.36 | 303.74 | 471.94 |
| 1672_5_weeks_ileum | 7.33 | 7.20 | 7.46 | 214.66 | 152.98 | 344.23 |
| 1573_5_weeks_jejunum | 3.89 | 3.78 | 4.01 | 301.47 | 247.64 | 398.11 |
| 1617_5_weeks_jejunum | 5.63 | 5.52 | 5.75 | 285.65 | 224.47 | 397.68 |
| 1621_5_weeks_jejunum | 2.92 | 2.84 | 3.00 | 336.00 | 293.00 | 410.43 |
| 1642_5_weeks_jejunum | 4.33 | 4.23 | 4.44 | 228.04 | 177.67 | 327.85 |
| Mock community control | 10.78 | 10.53 | 11.03 | 200.96 | 154.09 | 296.74 |

**Figure 1: UpSet graph – timepoints**

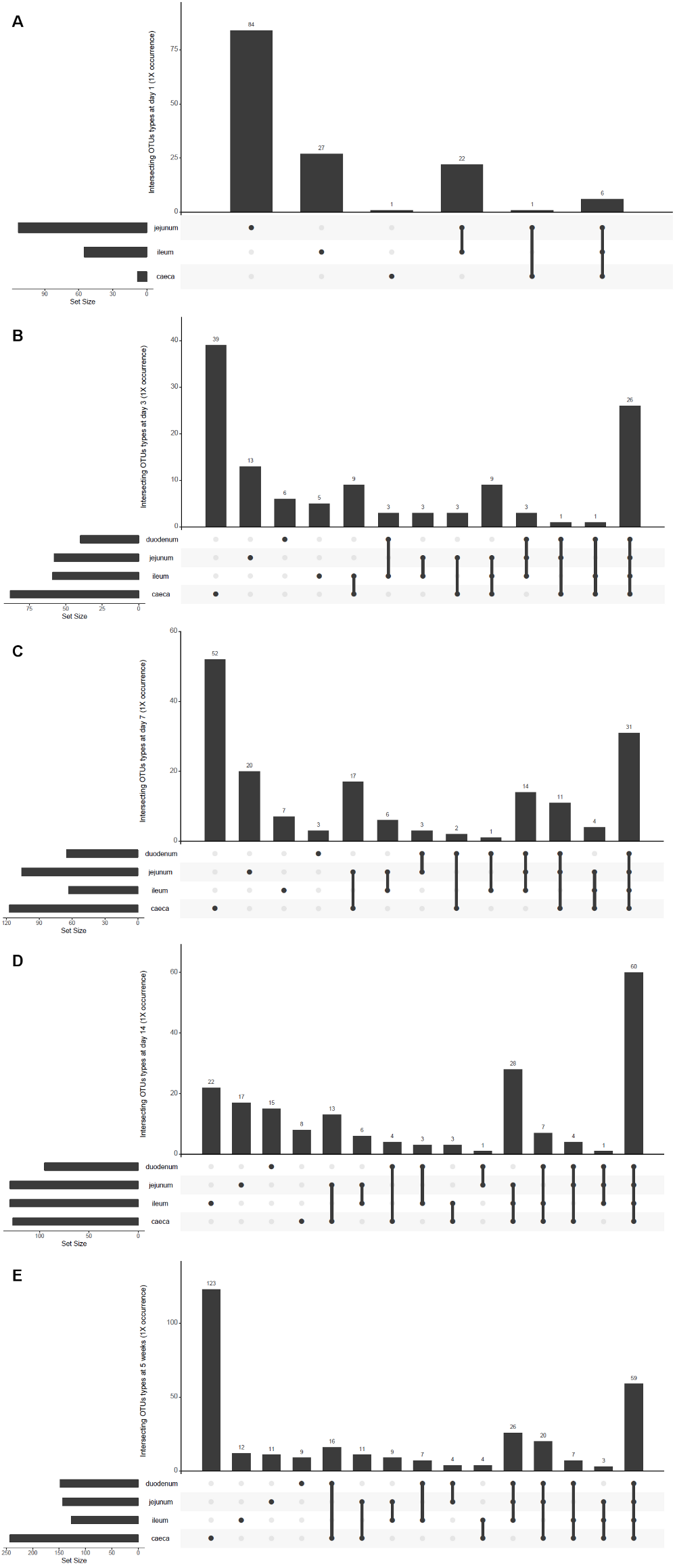

**UpSet graphs showing the number of shared OTUs between sample types at different timepoints (average 1 X coverage after subsampling to 10,000 reads). Duodenal samples from day 1 were not included due to scarcity of samples at this timepoint. A: day 1, B: day 3, C: day 7, D: day 14, E: 5 weeks.**

**Figure 2: UpSet graph – sample types**

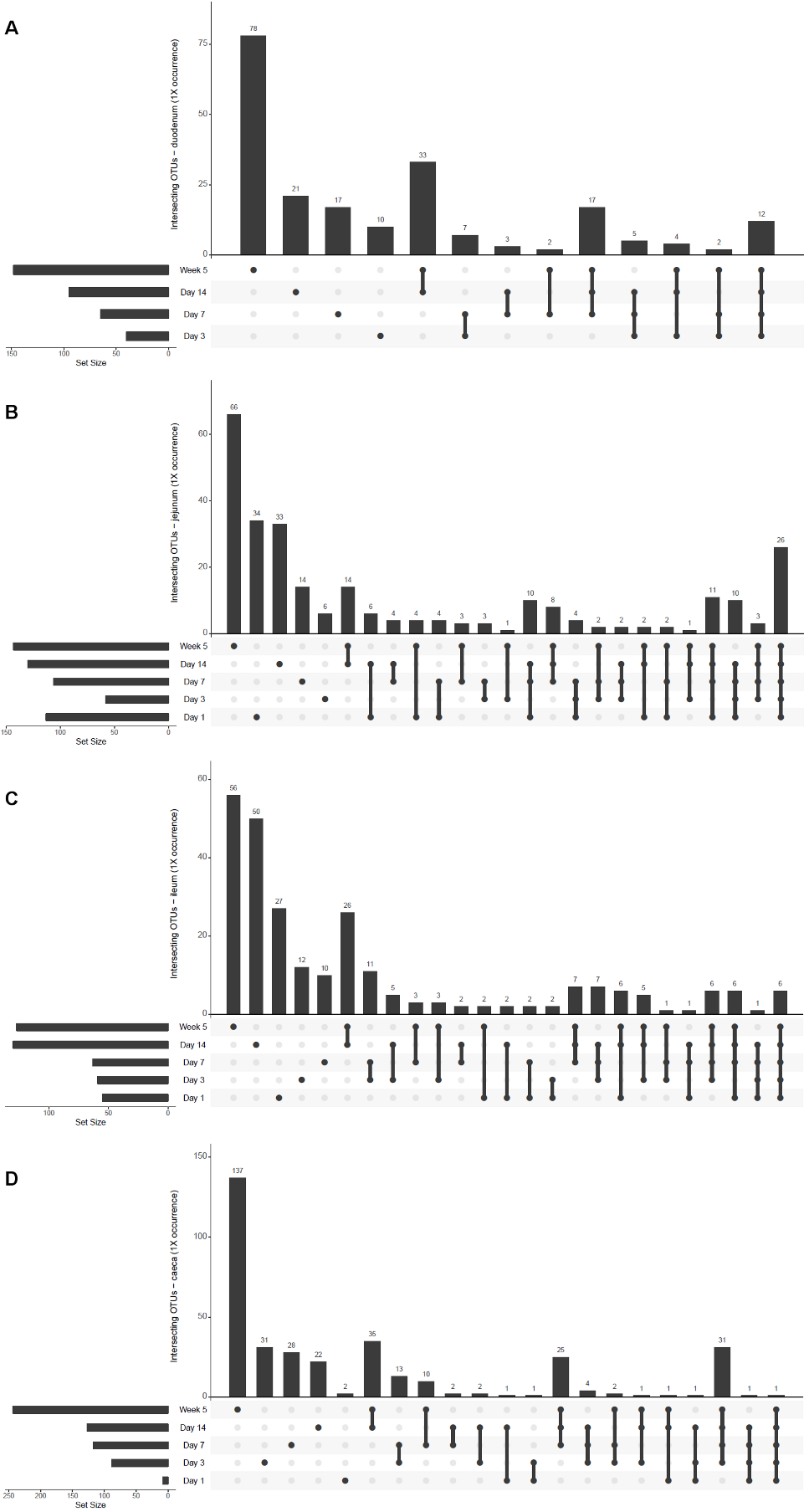

**UpSet graphs showing the number of shared OTUs between sample types at different timepoints (average 1 X coverage after subsampling to 10,000 reads). Duodenal samples from day 1 were not included due to scarcity of samples at this timepoint. A: duodenum, B: jejunum, C: ileum, D: caeca**
